# Distinct Core Glycan and O-Glycoform Utilization of SARS-CoV-2 Omicron Variant Spike Protein RBD Revealed by Top-Down Mass Spectrometry

**DOI:** 10.1101/2022.02.09.479776

**Authors:** David S. Roberts, Morgan Mann, Brad H. Li, Donguk Kim, Allan R. Brasier, Song Jin, Ying Ge

## Abstract

The SARS-CoV-2 Omicron (B.1.1.529) variant possesses numerous spike (S) mutations particularly in the S receptor-binding domain (S-RBD) that significantly improve transmissibility and evasion of neutralizing antibodies. But exactly how the mutations in the Omicron variant enhance viral escape from immunological protection remains to be understood. The S-RBD remains the principal target for neutralizing antibodies and therapeutics, thus new structural insights into the Omicron S-RBD and characterization of the post-translational glycosylation changes can inform rational design of vaccines and therapeutics. Here we report the molecular variations and O-glycoform changes of the Omicron S-RBD variant as compared to wild-type (WA1/2020) and Delta (B.1.617.2) variants using high-resolution top-down mass spectrometry (MS). A novel O-glycosite (Thr376) unique to the Omicron variant is identified. Moreover, we have directly quantified the Core 1 and Core 2 O-glycan structures and characterized the O-glycoform structural heterogeneity of the three variants. Our findings reveal high resolution detail of Omicron O-glycoforms and their utilization to provide direct molecular evidence of proteoform alterations in the Omicron variant which could shed light on how this variant escapes immunological protection.

## Introduction

The SARS-CoV-2 Omicron (B.1.1.529) variant has been classified by the World Health Organization (WHO) as a variant of concern (VOC) due to its significantly increased transmissibility, significant evasion of neutralizing antibodies from convalescents or vaccines, and higher risk of eluding testing.^1, 2^ Moreover, recent clinical data showed that this highly mutated Omicron variant causes higher rates of reinfection and rampant breakthrough infections with drastically different clinical outcomes as compared to wild-type (WT, WA1/2020) and Delta (B.1.617.2) variants, despite mRNA vaccine booster dose.^3^ Strikingly, the Omicron variant possess an alarming number of mutations (> 30), including 15 site mutations in the S receptor-binding domain (S-RBD) as compared to the WT strain. But how the mutations in the Omicron variant enhance viral escape from immunological protection remains to be understood.

Given that the S-RBD is the principal target for neutralizing antibodies and other therapeutics,^4^ and that glycosylation plays critical roles in host receptor ACE2 binding and function,^5-8^ it is crucial to decipher the glycoform changes of the Omicron S-RBD as compared to WT and Delta. Although S protein N-glycosylation has been characterized in detail^9, 10^ with ongoing efforts to understand the impact of N-glycosylation for vaccine development,^11^ characterization of O-glycosylation is challenging^12, 13^ due to the large microheterogeneity and structural diversity of O-glycans leading to multiple O-glycoforms.^14, 15^ To address these challenges, we have recently developed a hybrid top-down mass spectrometry (MS) approach^16^ for comprehensive characterization of O-glycoforms, along with other post-translational modifications (PTMs), to enable proteoform analysis^17, 18^ of complex glycoproteins. Because of the rapid mutation and spread of emerging SARS-CoV-2 variants, there is an urgent need for accurately distinguishing S-RBD variants and elucidating their post-translational glycosylation changes to bridge the knowledge gap between genomic changes and their clinical outcomes.

Here we report the first analysis of the sites and O-glycoform structure differences of the Omicron SARS-CoV-2 S-RBD variant compared to the WT and Delta variant, by top-down MS. Not only has a new O-glycan site been observed, but also top-down O-glycoform quantification revealed significant enhancement of Core 2 type O-glycan structures for Omicron, as compared to the WT or Delta variants.

## Results and Discussion

We used HEK293 expressed S-RBD protein arising from WT (WA1/2020), Delta (T478K), and Omicron (BA.1) variants for all the top-down MS analysis. The mutational differences inherent to the Delta and Omicron variants, as compared to the WT strain, are especially pronounced in their RBDs (Fig. 1A). To elucidate the molecular sequence and O-glycans of the various S-RBDs, we removed the N-glycans from the S-RBD using a PNGase F treatment (see Supplementary Information)^16^ to minimize the interference posed by N-glycan heterogeneity (Fig. 1B). N-glycans removal yielded a ∼10 kDa decrease in the molecular weight by SDS-PAGE as compared to the neat S-RBD. The resulting S-RBD O-glycoforms were resolved by an ultrahigh resolution 12 T Fourier transform ion cyclotron (FTICR)-MS (Fig.1C). Notably, the Omicron variant shows drastic differences in its O-glycosylation profile, as compared to the other variants (Fig. S1-S3). We utilized a timsTOF Pro capable of trapped ion mobility spectrometry (TIMS)-MS^19^ to separate and analyze the various S-RBD O-glycan structures for detailed glycoform characterization following the N-glycan removal (Fig. 2). To characterize the glycan structures and occupancy of the highly mutated Omicron variant, we performed specific isolation of the individual S-RBD O-glycoforms. Focusing on the most abundant O-glycoform (26+ charge state, 1069.4.3 *m/z*) as a specific example, collisionally activated dissociation (CAD) fragment ions were analyzed in targeted protein analysis mode using MASH Explorer^20^. The top-down MS/MS spectra were obtained along with ion mobility separation of the various O-glycan structures to overcome the microheterogeneity inherent to O-glycans analysis (Fig. 2B). Soft TIMS cell activation parameters enabled detailed neutral loss mapping of the isolated S-RBD proteoform and revealed a Core 1 (Galβ1‐3GalNAc‐Ser/Thr) O-glycan with a GalNAcGal(NeuAc)_2_ structure (Fig. 2C). Taking advantage of the TIMS separation, we could accurately assign the specific isolated precursor to a single parent O-glycan structure, thereby overcoming mass degeneracy inherent to glycan assignment.^21^ Moreover, this TIMS-MS approach revealed multiple S-RBD glycoforms with Core 1 and Core 2 (GlcNAcβ1‐6(Galβ1-3)GalNAc‐Ser/Thr) O-glycan structures across the three S-RBD variants (Fig. S4). Interestingly, the Omicron variant S-RBD presented different O-glycan patterns as compared to the WT and Delta variants.

**Figure 1.**
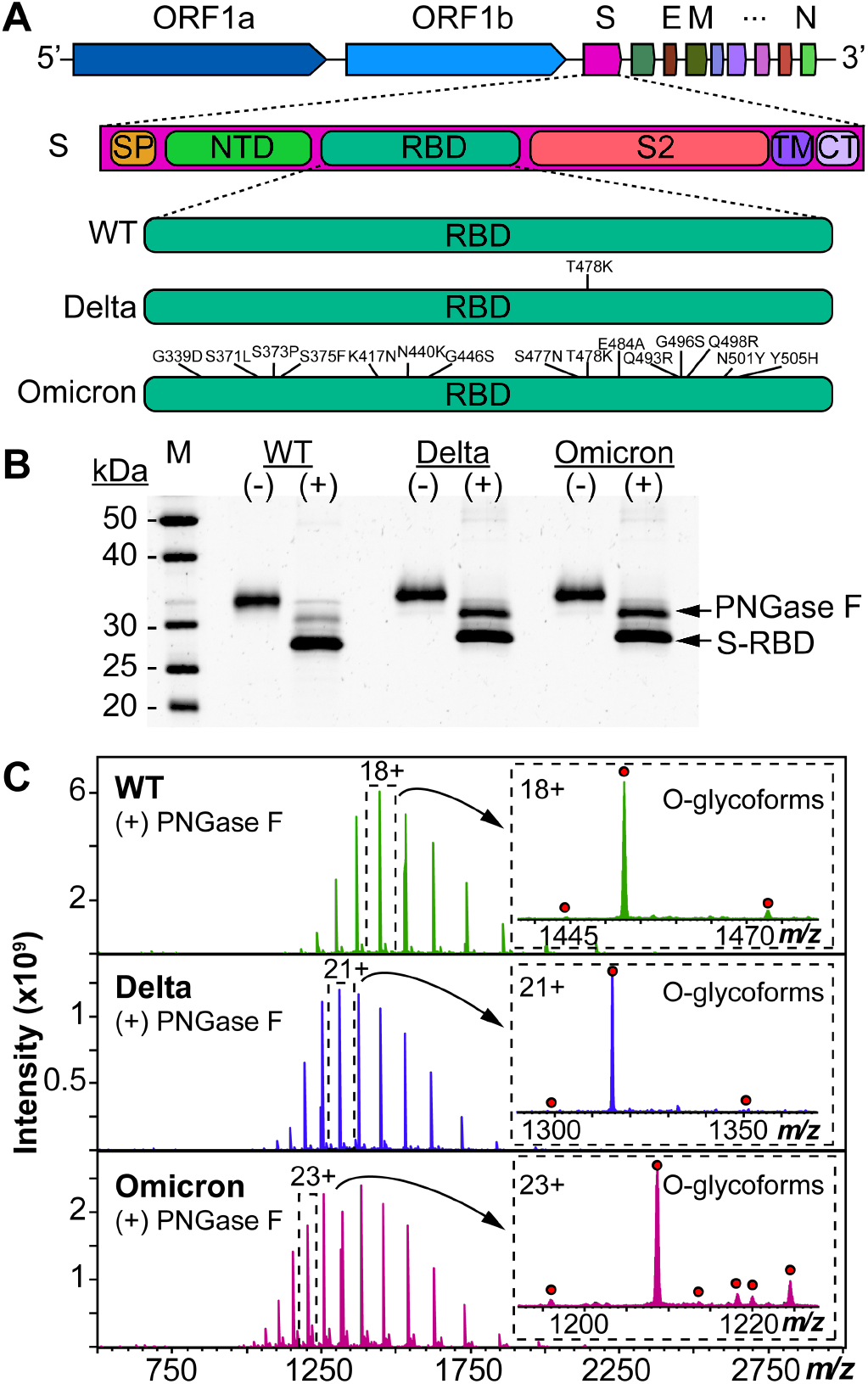
Protein mutational mapping and high-resolution top-down MS of S-RBD arising from the WT, Delta, and Omicron variants. (A) Architecture of SARS-CoV-2 genome and illustration of the protein sequence changes for the S-RBD variants. (B) SDS-PAGE of the S-RBD variants before (−) and after (+) PNGase F treatment. Gel lanes were equally loaded (300 ng), and gel staining was visualized by SYPRO Ruby. (C) Raw MS1 of intact S-RBD proteoforms collected on the 12 T FTICR-MS after PNGase F treatment for the WT, Delta, and Omicron variants. All identified O-glycoforms are annotated in the inset with a red circle.

**Figure 2.**
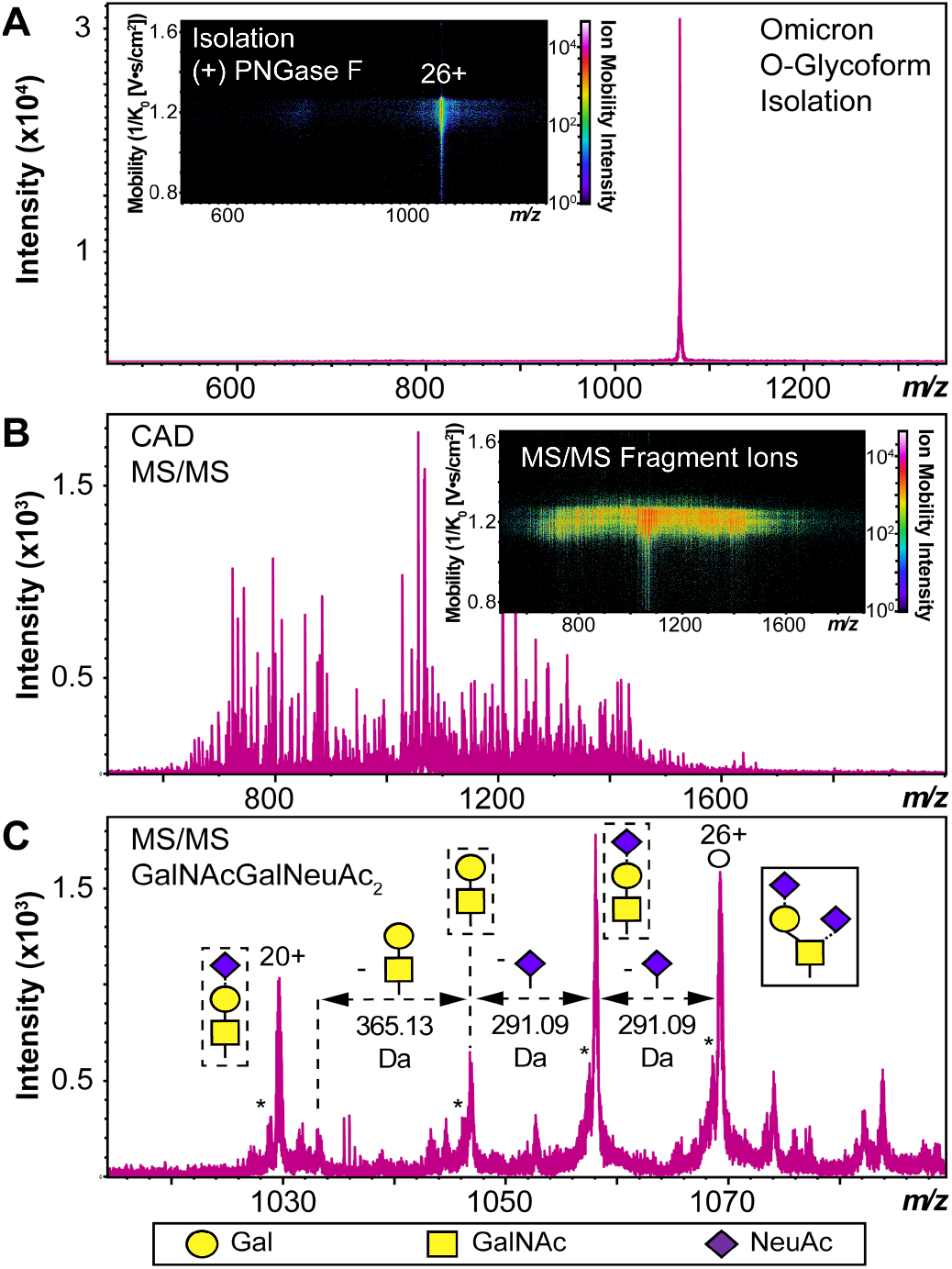
S-RBD O-glycoform analysis by TIMS-MS/MS. (A) Illustration of the isolation of a specific Omicron S-RBD glycoform (z = 26+, centered at 1069.4 m/z) after PNGase F treatment using TIMS-MS. (Inset) Associated ion mobility heat map after isolation of the precursor. (B) Top-down MS/MS of Omicron S-RBD O-glycoform following CAD fragmentation of the isolated proteoform. A collision cell voltage of 22 eV was applied for this specific example. (C) Neutral loss O-glycan mapping of the Omicron S-RBD proteoform. Glycoform characterization reveals the specific S-RBD proteoform to have a Core 1 type GalNAcGalNeuAc2 glycan structure. Assignments of the glycan structures are marked in the spectrum with the legend shown on the bottom. The hollow circle represents the 26+ charge state precursor ion corresponding to the 1069.4 m/z isolation. An asterisk “*” denotes an oxonium ion loss.

We then further characterized the S-RBD WT, Delta, and Omicron O-glycosylation patterns to reveal all the structural O-glycoform alterations between the variants (Fig. 3). Interestingly, we observed major O-glycan microheterogeneity changes in Omicron, as compared to the WT or Delta variants. In particular, we found significantly enhanced Core 2 O-glycan structure abundances for Omicron with pronounced expression for multiply sialylated GalNAc(GalNeuAc)(GlcNAcGalNeuAc) and fucosylated GalNAc(GalNeuAc)(GlcNAcGalFuc) structures. The striking molecular abundance differences observed in Omicron as compared to the WT or Delta variants are summarized in Table 1. The relative abundance of Core 1 to Core 2 S-RBD O-glycan structures for the Omicron variant was roughly 71:29, with the Core 1 GalNAcGal(NeuAc)_2_ being the most abundant O-glycoform (∼69% relative abundance). Interestingly, the WT and Delta variants show a strong bias toward Core 1 type O-glycan structures, with more than 80% of its O-glycoform abundance corresponding to the Core 1 GalNAcGal(NeuAc)_2_ structure. The Omicron variant was found to possess a Core 2 type GalNAc(GalNeuAc)(GlcNAcGalFuc) structure that accounts for more than 13% of its total O-glycoform composition (Fig. 3 and S5). Moreover, we characterized the abundant (10 %) Core 2 GalNAc(GalNeuAc)(GlcNAcGalNeuAc) structure for the Omicron variant (Fig. 3 and S6). These particular Core 2 structures were also found in the WT and Delta S-RBD variants but at much lower (5-7%) relative abundances. The high-resolution intact S-RBD glycoform characterization shown in Fig. 3 demonstrates the distinct advantages of this top-down MS approach over glycopeptide-based bottom-up MS approaches.^22, 23^

**Figure 3.**
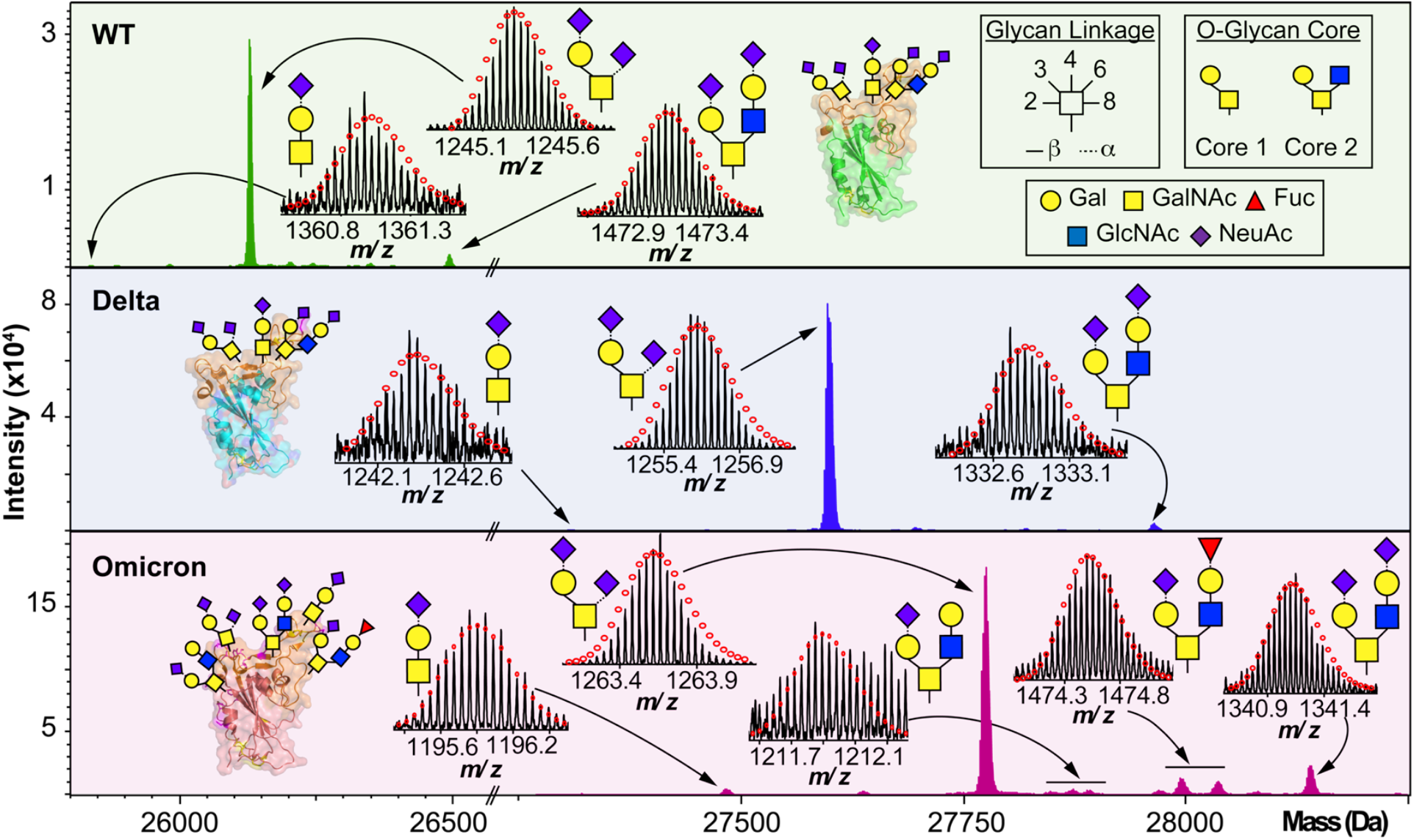
O-glycoform characterization of the S-RBD variants. Deconvoluted MS of the S-RBD proteoforms are shown with all major O-glycan assignments for the WT (green), Delta (blue), and Omicron (pink) variants. Insets show the isotopically and baseline resolved S-RBD proteoform MS spectra allowing for accurate determination of individual intact glycoforms. MS data were collected on the 12 T FTICR-MS. All individual ion assignments are within 1 ppm from the theoretical mass and theoretical isotopic distributions are indicated by the red dots. Assignments of the glycan structures are marked in the spectrum with the legend shown as an inset. PDB: 6M0J, 7WBQ, 7WBP.

**Table 1.** Summary of the S-RBD variants O-glycoform relative abundances.

| O-glycoform structure | Cr.<sup>[a]</sup> | WT | Delta | Omicron |
| --- | --- | --- | --- | --- |
|  |  | (%) | (%) | (%) |
| GalNAcGalNeuAc | 1 | 1.3 | 1.4 | 2.3 |
| GalNAcGal(NeuAc)<sub>2</sub> | 1 | 80.9 | 86.2 | 69.0 |
| GalNAc(GalNeuAc)(GlcNAcGal)<sup>[b]</sup> | 2 | 5.8 | 3.8 | 5.4 |
| GalNAc(GalNeuAc)(GlcNAcGalFuc)<sup>[b]</sup> | 2 | 5.2 | 3.2 | 13.2 |
| GalNAc(GalNeuAc)(GlcNAcGalNeuAc) | 2 | 6.8 | 5.3 | 10.0 |

[a] O-glycan core type. [b] Combined abundance including O-glycoforms with a N-terminal acetylation (WT, Delta) or N-terminal trimethylation (Omicron).

We then further investigated the glycosites and their microheterogeneity between the S-RBD variants. Fig. 4 shows a specific example of top-down MS/MS O-glycosite determination using the highly abundant Core 1 GalNAcGal(NeuAc)_2_ structure. Detailed top-down MS/MS analysis of the S-RBD O-glycoforms revealed the presence of a new O-glycosite (Thr376) unique to the Omicron variant (Fig. 4A). Fascinatingly, all detected S-RBD O-glycans for the WT and Delta variants were confidently assigned solely to Thr323 (Fig. 4B and 4C), which agrees with previous studies on WT S O-glycosylation.^24^ Representative CAD fragment ions for the WT (*b*_*7*_^1+^ and *b* _173_^11+^ and Delta (*b*_*7*_ ^1+^ and *b*_233_ ^12^) confidently localized the O-glycosite to Thr323 (Fig. 4b-c and S7). Given the smaller number of mutations present on Delta as compared to Omicron, it is not surprising that the O-glcyosite Thr323 was conserved between Delta and WT variants. On the other hand, the Omicron variant yielded both the familiar Thr323 O-glycosite (*b*_5_^1+^ and *b*_18_^2+^) and a new Thr376 O-glycosite (*b*_60_ ^12+^ and *b*_52_ ^5+^) corresponding to the GalNAcGal(NeuAc) O-glycoform that is simultaneously occupied (Fig. 4D). This Thr376 O-glycosite is conveniently n + 3 adjacent to a proline at residue 373, which is consistent with previous reports of increased O-glycosylation frequency near proline.^25^ This particular Pro373 is a site-specific mutation unique to the Omicron variant and likely is the reason for this new O-glycosite. We note that the site occupancy of the Thr376 site is low(< 30%) relative to the Thr323 and was only confidently assigned for the abundant GalNAcGal(NeuAc)_2_ O-glycoform, although we suspect other Core 2 O-glycoforms may also possess the Thr376 modification. It should be noted that recombinant fragments of S protein may show differential glycosylation when the S-RBD is expressed as a monomer, therefore care is needed when assigning O-glycans between different protein sources.^26^ Moreover, although the S-RBD O-glycoforms assigned for the variants are specific to HEK293 derived S-RBD, the HEK293 expression model has been shown to reflect glycosylation sites expected for the viron.^27-29^

**Figure 4.**
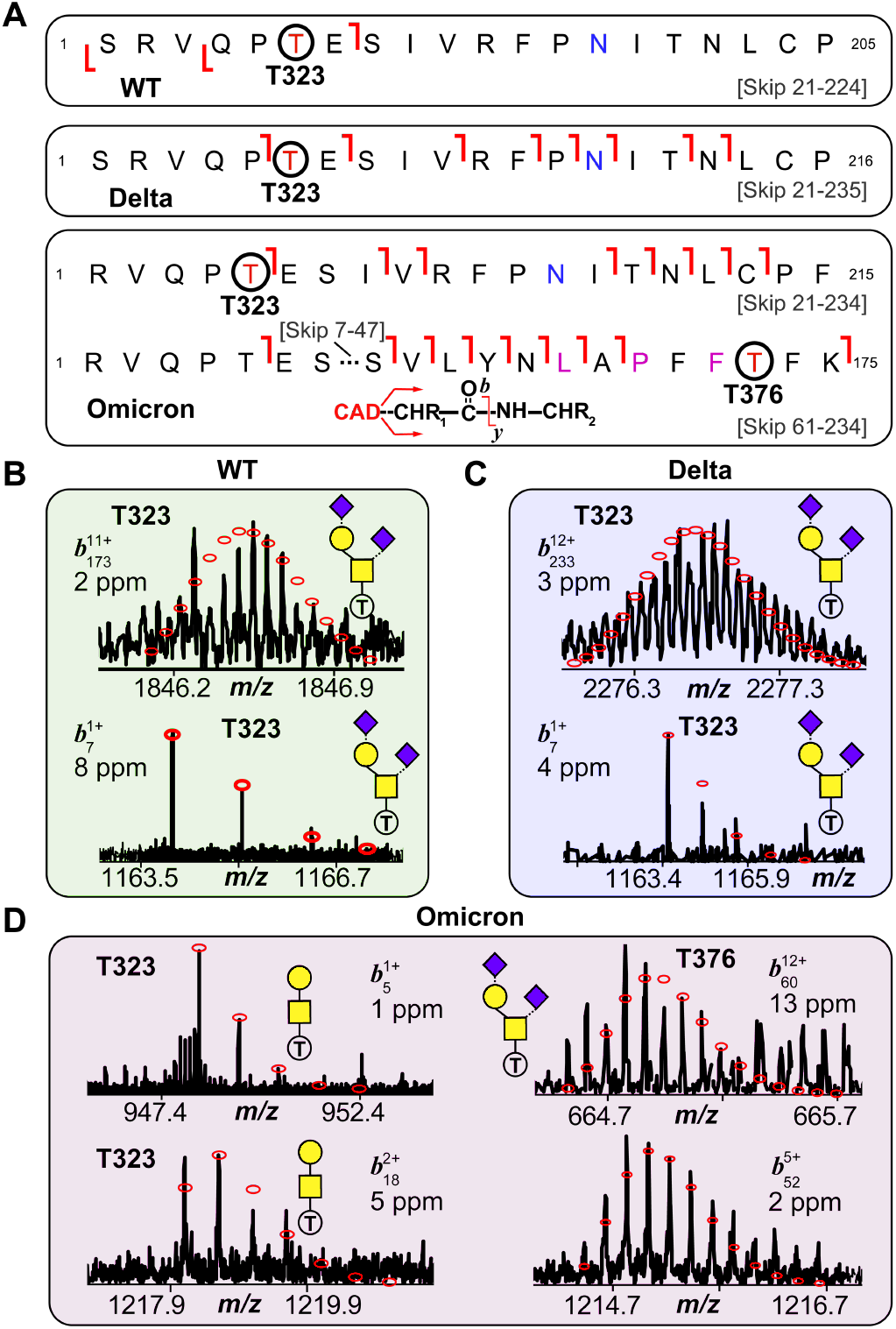
S-RBD O-glycosite localization by top-down TIMS-MS/MS. (A) Fragmentation mapping of the Core 1 type GalNAcGal(NeuAc)2 glycan corresponding to the WT, Delta, and Omicron S-RBD variants. Amino acid sequence (Arg319-Phe541) was based on the entry name P0DTC2 (SPIKE_SARS2) obtained from the UniProtKB sequence database with appropriate amino acid substitutions made for the mutated Delta and Omicron variants. Cleavage of signal peptide tPA following cell expression results in the N-terminal Ser for the WT and Delta variants. The blue N (Asn) denotes deamidation following PNGase F treatment. Specific Omicron residue mutations are denoted with a pink residue. (B-D) Representative top-down MS/MS CAD fragment ions of intact WT (b71+ and b17311+) (B), Delta (b71+ and b22312+) (C), and Omicron (b51+, b182+, b71+, b17311+) (D) variants corresponding to the GalNAcGal(NeuAc)2 O-glycoform shown in (A). WT and Delta variants showed complete O-glycosite occupation at Thr323. The Omicron variant was found to possess both Thr323 (left side; b51+and b182+) and Thr376 (right side; b71+, and b17311+) as O-glycosites corresponding to the same Core 1 O-glycan structure. Theoretical isotopic distributions are indicated by the red dots and mass accuracy errors are listed for each fragment ion.

## Conclusion

In summary, we report the first analysis of the O-glycoform structural heterogeneity of the S-RBD found in the SARS-CoV-2 Omicron variant. We observed significant enhancement in the utilization of Core 2 type O-glycoforms for the Omicron variant as compared to WT or Delta. Moreover, we identified and characterized a novel Thr376 O-glycosite unique to Omicron S-RBD. This top-down MS approach is complimentary to traditional structural methods such as X-ray crystallography and cryoEM, which are not amenable for direct glycan structural analysis due to the inherent flexibility and heterogeneity of oligosaccharides,^30, 31^ and provides unmatched resolution for the characterization of SARS-CoV-2 S-RBD proteoforms. Importantly, this top-down MS approach can be leveraged to resolve the protein level changes of S mutations in emerging SARS-CoV-2 variants to understand changes in their structure-function. Our findings bridge the knowledge gap between S variant genomic alterations and final clinical outcomes with detailed proteoform information, which could shed new light on how Omicron escapes immunological protection.

## Acknowledgement

This research is supported by NIH R01 GM117058 (to S.J. and Y.G.). D.S.R. acknowledges the support from the American Heart Association Predoctoral Fellowship Grant No. 832615/David S. Roberts/2021. Y.G. would like to acknowledge NIH R01 GM125085, R01 HL096971, and S10 OD018475. We also thank Michael Greig, Guillaume Tremintin, Conor Mullins, Gary Kruppa, Paul Speir, and Rohan Thakur of Bruker Daltonics for their helpful discussion and provision of the Bruker timsTOF Pro used in this work.

## Notes

The authors declare no competing financial interest.

