## Supplementary Materials for "Distinct Core Glycan and O-Glycoform Utilization of SARS-CoV-2 Omicron Variant Spike Protein RBD Revealed by Top-Down Mass Spectrometry"

**Table of Contents** **Page**


**Materials and Methods** Pages 3-5

**Figure S1:** MS illustration of intact S-RBD WT variant Page 6

**Figure S2:** MS illustration of intact S-RBD Delta variant Page 7

**Figure S3:** MS illustration of intact S-RBD Omicron variant Page 8

**Figure S4:** S-RBD variants O-glycoform proteoform summary Page 9

**Figure S5:** MS/MS characterization of Omicron GalNAc(GalNeuAc)(GlcNAcGalFuc) Page 10

**Figure S6:** MS/MS characterization of Omicron GalNAc(GalNeuAc)(GlcNAcGalNeuAc) Page 11

**Figure S7:** O-glycosylation site characterization for the Omicron S-RBD Page 12

**References** Page 13

### Materials and Methods

**Materials and Reagents.** All chemicals and reagents were purchased from MilliporeSigma (Burlington, MA, USA) and used as received without further purification unless otherwise noted. Aqueous solutions were made in nanopure deionized water ($H_2O$) from Milli-Q® water (MilliporeSigma). Ammonium acetate (AA) was purchased from Fisher Scientific (Fair Lawn, NJ). Bradford protein assay reagent was purchased from Bio-Rad (Hercules, CA, USA). 12.5% gel (10 or 15 comb well, 10.0 cm × 10.0 cm, 1.0 mm thick) for SDS-Polyacrylamide Gel Electrophoresis (SDS-PAGE) was home-made. Thermo Scientific™ Cimarec+™ stirring hotplate were purchased from ThermoFisher Scientific. Amicon, 0.5 mL cellulose ultra centrifugal filters with a molecular-weight cutoff (MWCO) of 10 kDa were purchased from MilliporeSigma. The recombinant SARS-CoV-2 spike regional-binding domain protein expressed in the HEK 293 cell line was purchased from Sino Biological Inc. (cat. 40592-VNAH, 40592-V08H91, and 40592-V08H121).

SARS-CoV-2 Spike RBD WT (WA1/2020):
SRVQPTESIVRFPNITNLCPFGEVFNATRFASVYAWNRKRISNCVADYSVLYNSASFSTF
KCYGVSPTKLNDLCFTNVYADSFVIRGDEVRQIAPGQTGKIADYNYKLPDDFTGCVIAW
NSNNLDSKVGGNYNYLYRLFRKSNLKPFERDISTEIYQAGSTPCNGVEGFNCYFPLQSY
GFQPTNGVGYQPYRVVVLSFELLHAPATVCGPKKSTNLVKNKCVNF

SARS-CoV-2 Spike RBD Delta (T478K):
SRVQPTESIVRFPNITNLCPFGEVFNATRFASVYAWNRKRISNCVADYSVLYNSASFST
FKCYGVSPTKLNDLCFTNVYADSFVIRGDEVRQIAPGQTGKIADYNYKLPDDFTGCVIA
WNSNNLDSKVGGNYNYLYRLFRKSNLKPFERDISTEIYQAGSKPCNGVEGFNCYFPLQ
SYGFQPTNGVGYQPYRVVVLSFELLHAPATVCGPKKSTNLVKNKCVNFAHHHHHHHH
HH

SARS-CoV-2 Spike RBD Omicron (BA.1):
RVQPTESIVRFPNITNLCPFDEVFNATRFASVYAWNRKRISNCVADYSVLYNLAPFFTF
KCYGVSPTKLNDLCFTNVYADSFVIRGDEVRQIAPGQTGNIADYNYKLPDDFTGCVIAW
NSNKLDSKVSGNYNYLYRLFRKSNLKPFERDISTEIYQAGNKPCNGVAGFNCYFPLRS
YSFRPTYGVGHQPYRVVVLSFELLHAPATVCGPKKSTNLVKNKCVNFAHHHHHHHHH
H

**Complete N-glycan removal for O-glycan protein analysis.** Peptide-N-glycosidase F (PNGase F) (New England Biolabs Inc., Cat. P0704S) was used for complete removal of N-linked oligosaccharides. Briefly, 20 µg of protein sample was buffer exchanging into 150 mM AA (pH = 7.4) solution by washing the sample five times through a 10 kDa Amicon ultra centrifugal filters (MilliporeSigma, Burlington, MA, USA). PNGase F (5 U) was then added to the protein solution and samples were incubated at 37 °C for 12 h on a Thermo Scientific™ Cimarec+™ stirring hotplate.

**Sample preparation.** Protein samples were reduced using 20 mM tris(2-carboxyethyl)phosphine hydrochloride (TCEP), and followed by buffer exchanging into 0.1 % formic acid solution by washing the sample five times through a 10 kDa Amicon ultra centrifugal filters (MilliporeSigma, Burlington, MA, USA). The protein sample was then diluted to 20 µM in 50:50:0.1 (acetonitrile:

water: formic acid) prior to top-down MS analysis. Protein samples subjected to N-glycan removal were prepared similarly following PNGase F reaction.

**FTICR-MS top-down analysis.** Intact protein samples were analyzed by nano-electrospray ionization via direct infusion using a TriVersa Nanomate system (Advion BioSciences, Ithaca, NY, USA) coupled to a solariX XR 12-Tesla Fourier Transform Ion Cyclotron Resonance mass spectrometer[1, 2] (FTICR-MS, Bruker Daltonics, Billerica, MA). For the nano-electrospray ionization source using a TriVersa Nanomate, the desolvating gas pressure was set at 0.45 PSI and the voltage was set to 1.2 to 1.6 kV versus the inlet of the mass spectrometer. Source dry gas flow rate was set to 4 L/min at 124 °C. For the source optics, the capillary exit, deflector plate, funnel 1, skimmer voltage, funnel RF amplitude, octopole frequency, octopole RF amplitude, collision cell RF frequency, and collision cell RF amplitude were optimized at 190 V, 200 V, 100 V, 35 V, 250 Vpp, 2 MHz, 490 Vpp, 2MHz, and 2000 Vpp, respectively. Mass spectra were acquired with an acquisition size of 4M-words of data (with a resolution of 530,000 at 400 *m/z*), in the mass range between 200-4000 *m/z*, and a minimum of 250 scans were accumulated for each sample. Ions were accumulated in the collision cell for 0.100 s, and a time of flight of 1.500 ms was used prior to their transfer to the ICR cell. For collisionally activated dissociation (CAD) tandem MS (MS/MS) experiments, the collision energy was varied from 10 to 30V, ion accumulation was optimized to 400 ms, and acquisition size was 4M-words of data. Tandem mass spectra were output from the DataAnalysis 4.3 (Bruker Daltonics) software and analyzed using MASH Explorer.[3]

**TIMS-MS top-down analysis.** A timsTOF Pro mass spectrometer (Bruker Daltonics, Bremen, Germany) was coupled to a Bruker nanoElute LC system (Bruker Daltonics, Bremen, Germany). Samples were directly infused using the nanoElute, injecting 5 μL of protein sample with a flow rate of 1 μL/min. For the MS inlet, the endplate offset and capillary voltage were set to 500 V and 3800 V, respectively. The nebulizer gas pressure ($N_2$) was set to 1.0 bar, with a dry gas flow rate of 5 L/min at 200 °C. The tunnel out, tunnel in, and TOF vacuum pressures were set to 0.8584 mBar, 2.628 mBar, and 1.752E-07 mBar. To calibrate the MS and trapped ion mobility spectrometry (TIMS) device, Agilent tune mix was directly infused to provide species of known mass and reduced mobility.[37] For MS calibration, the MS resolution for the most abundant calibrant signal, 1222 *m/*z, was 72,000. Calibrant points at 922, 1222, and 1522 *m/z* were used for TIMS calibration. The TIMS resolution for the most abundant calibrant signal, 1222 *m/*z, was 84.2 CCS/ΔCCS. IMS tunnel voltage deltas were optimized at -20 V, -120 V, 30 V, 100 V, 0 V and 100 V for Δ1 to Δ6, respectively. TIMS funnel 1 RF was set to 280 Vpp, and TIMS collision cell energy was set to 50 V. An IMS imeX accumulation time of 50 ms and cycle ramp time of 200 ms was found to yield optimal resolving power. The TIMS accumulation time was locked to the mobility range (typically 0.75 to 1.65 $1/K_0$). In the MS transfer optics the funnel 1 RF, funnel 2 RF, deflection delta, in-source collision induced dissociation (isCID) energy, multipole RF, and quadrupole ion energy were optimized at, 350 Vpp, 5 Vpp, 70 V, 50 eV, 200 Vpp, and 2 eV, respectively. For $MS^1$ spectral collection, the quadrupole low mass was set to 500 *m/z* with a scan range of 200 to 3000 *m/z.* Collision energy was set to 8 eV, with 2600 Vpp collision cell RF, a 110 μs transfer time, and a pre pulse storage time of 22 μs. For $MS^2$ spectral collection the collision energy was varied from 10 to 30V, the quadrupole low mass was set to 200 *m/z* with a scan range of 200 to 3000 *m/z.* The collision cell RF was set to 2600 Vpp, a 110 μs transfer time, and a pre pulse storage time of 22 μs were used. Mass spectra were output from the DataAnalysis 5.3 (Bruker Daltonics) software and analyzed using MASH Explorer.[3]

**Data Analysis.** All data were processed and analyzed using Compass DataAnalysis 4.3/5.3 and MASH Explorer.[3] Maximum Entropy algorithm (Bruker Daltonics) was used to deconvolute all mass spectra with the resolution set to 80,000 for the timsTOF Pro or with instrument peak width set to 0.05 for the 12T FTICR. Sophisticated Numerical Annotation Procedure (SNAP) peak-picking algorithm (quality factor: 0.4; signal-to-noise ratio (S/N): 3.0; intensity threshold: 500) was applied to determine the monoisotopic mass of all detected ions. The relative abundance for each protein isoform was determined using DataAnalysis. To quantify protein modifications, the relative abundances of specific modifications were calculated as their corresponding percentages among all the detected protein forms in the deconvoluted averaged mass. MS/MS data were output from the DataAnalysis software and analyzed using MASH Explorer[3] for proteoform identification and sequence mapping. All the program-processed data were manually validated. For peak picking, the sophisticated numerical annotation procedure (SNAP) algorithm from Bruker DataAnalysis 5.3 was used with a quality threshold of 0.5 and an S/N lower threshold of 3. All fragment ions were manually validated using MASH Explorer. Peak extraction was performed using a signal-to-noise ratio of 3 and a minimum fit of 60%, and all peaks were subjected to manual validation. All identifications were made with satisfactory numbers of assigned fragment (>10), and a 25-ppm mass tolerance was used to match the experimental fragment ions to the calculated fragment ions based on amino acid sequence. For ion mobility analysis by the timsTOF Pro, the collisional cross section (CCS) in Å$^2$ for a species of interest was determined via the Mason-Schamps equation (Equation 1):

$$CCS = \frac{3}{16}\sqrt{\frac{2\pi}{\mu k_b T}}\frac{ze}{N_0 K_0} \tag{1}$$

Where $\mu$ is the reduced mass of the ion-gas pair ($\mu = \frac{mM}{(m+M)}$, where m and M are the ion and gas particle masses), $k_b$ is Boltzmann's constant, $T$ is the drift region temperature, $z$ is the ionic charge, $e$ is the charge of an electron, $N_0$ is the buffer gas density, and $K_0$ is the reduced mobility. Equation 1 was selected to agree with previously published CCS calculations.[4-6] Theoretical CCS values were determined using the IMPACT method.[7]

**Supplementary Figures**

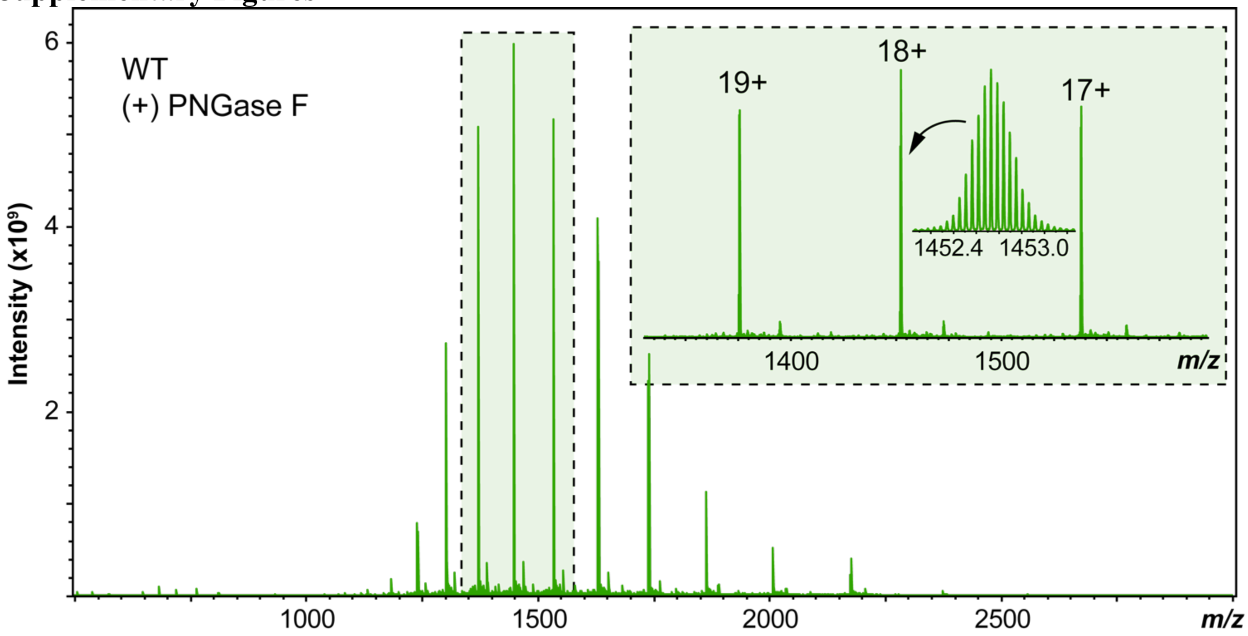


**Figure S1.** FTICR-MS analysis of intact S-RBD WT variant after PNGase F treatment showing baseline isotopic resolution of specific S-RBD proteoforms. The top three most abundant charge states (19+, 18+, and 17+) are highlighted in the inset, along with a zoom-in of the most abundant protoeoform illustrating the isotopically resolved spectrum.

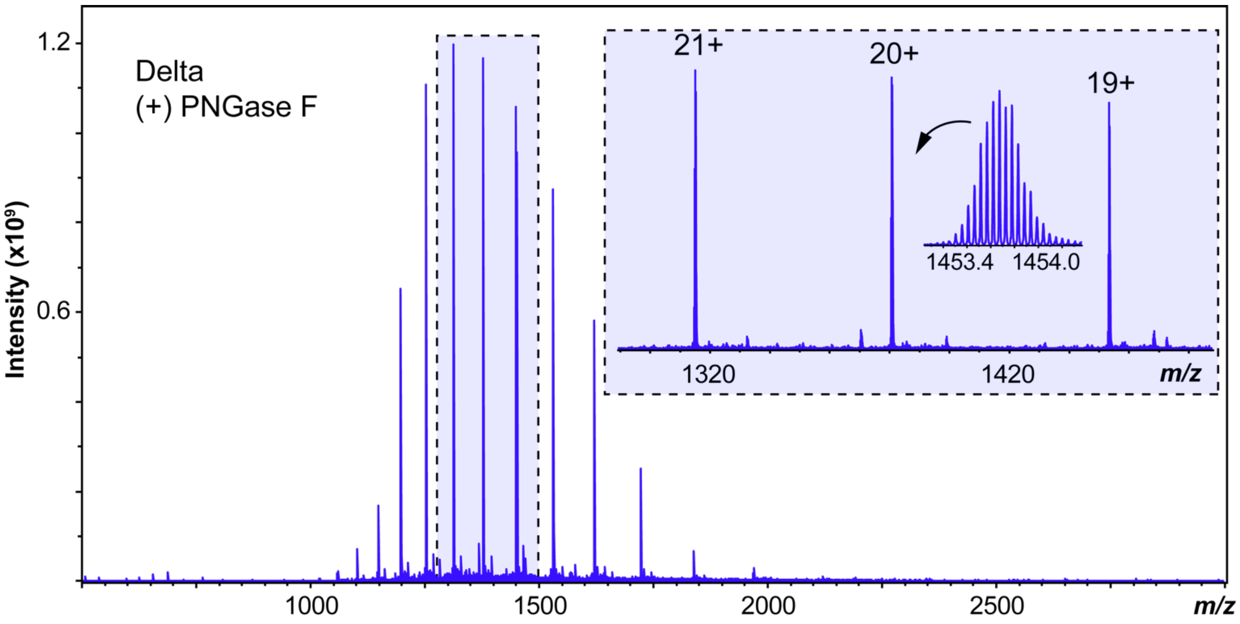


**Figure S2.** FTICR-MS analysis of intact S-RBD Delta variant after PNGase F treatment showing baseline isotopic resolution of specific S-RBD proteoforms. The top three most abundant charge states (21+, 20+, and 19+) are highlighted in the inset, along with a zoom-in of the most abundant proteoform illustrating the isotopically resolved spectrum.

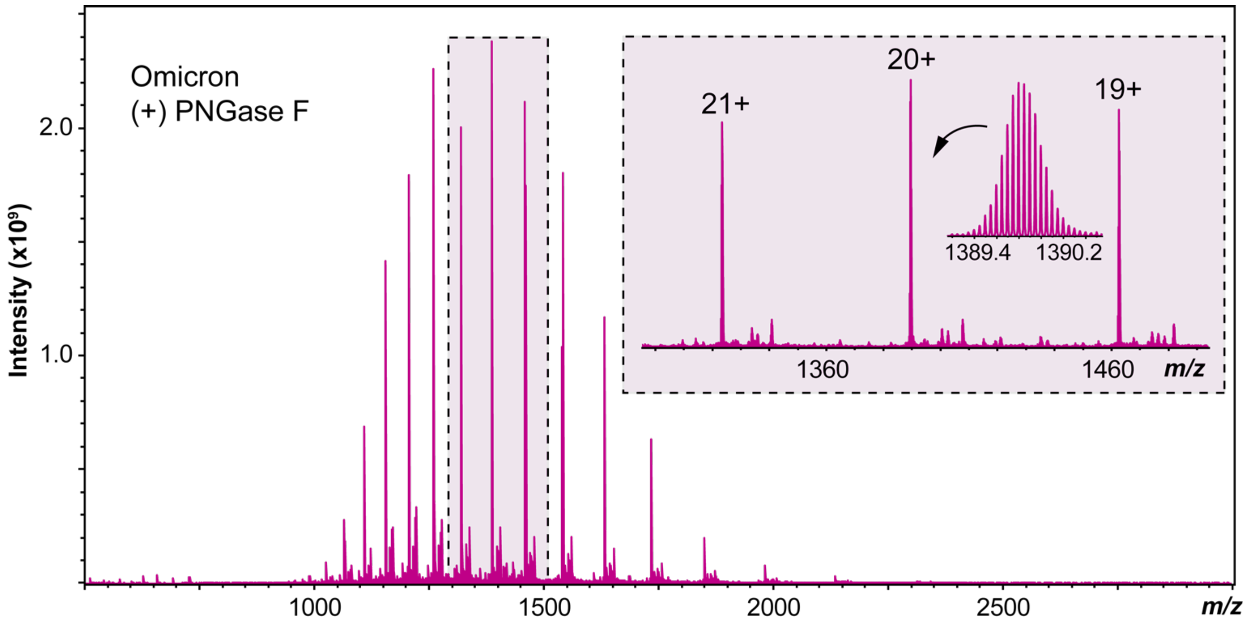


**Figure S3.** FTICR-MS analysis of intact S-RBD Delta variant after PNGase F treatment showing baseline isotopic resolution of specific S-RBD proteoforms. The top three most abundant charge states (21+, 20+, and 19+) are highlighted in the inset, along with a zoom-in of the most abundant proteoform illustrating the isotopically resolved spectrum.

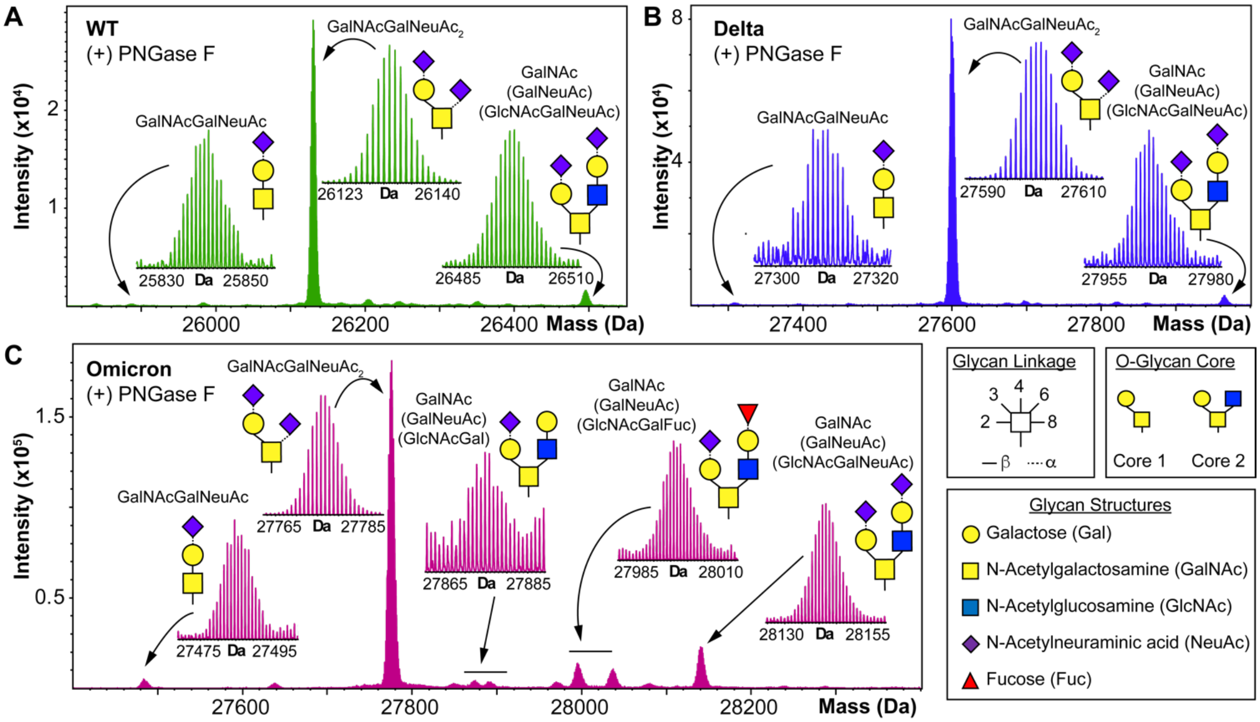


**Figure S4.** S-RBD O-glycoform analysis by high-resolution top-down MS for the WT, Delta, and Omicron variants. (A-C) Deconvoluted MS of the S-RBD O-glycoforms showing assignments for all identified O-glycoforms revealed by top-down MS for the (A) WT, (B) Delta, and (C) Omicron variants. The insets show a zoom-in of the individual O-glycoforms, illustrating the baseline isotopically resolved peaks. Assignments of the glycan structures are marked on the insets and are defined by the legend shown on the right side.

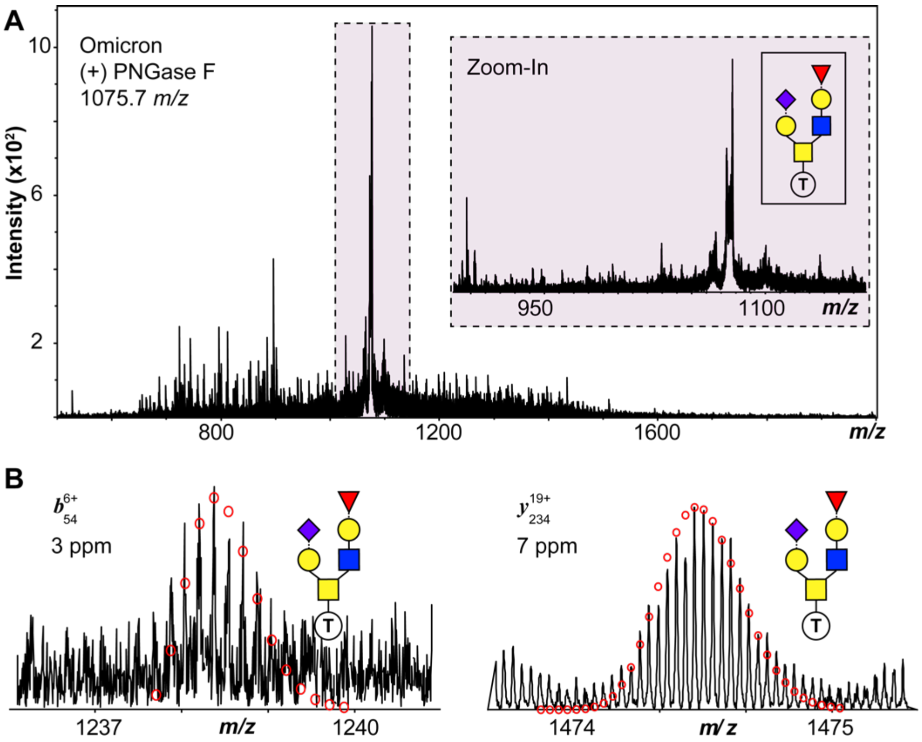


**Figure S5.** (A) Top-down MS/MS analysis of Omicron S-RBD proteoform isolated from quadrupole window centered at 1075.7 *m/z*, corresponding to the MS data shown in Figure 3. Glycoform characterization reveals the specific S-RBD proteoform to have Core 2 type GalNAc(GalNeuAc)(GlcNAcGalFuc) glycan. Inset shows a zoom-in region of the isolated O-glycoform. (B) Representative CAD fragment ions ($b_{54}^{6+}$, $y_{234}^{19+}$) obtained from S-RBD with glycosite at Thr323.

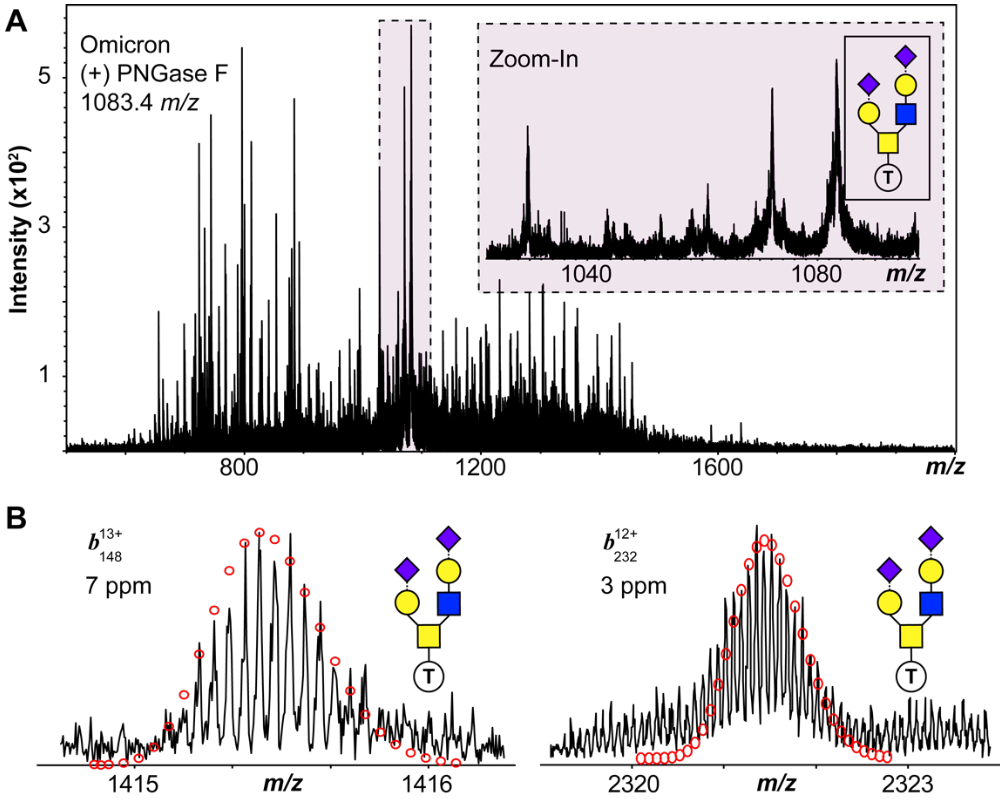


**Figure S6.** (A) Top-down MS/MS analysis of Omicron S-RBD proteoform isolated from quadrupole window centered at 1083.4 *m/z*, corresponding to the MS data shown in Figure 3. Glycoform characterization reveals the specific S-RBD proteoform to have Core 2 type GalNAc(GalNeuAc)(GlcNAcGalNeuAc) glycan. Inset shows a zoom-in region of the isolated O-glycoform. (B) Representative CAD fragment ions ($b_{148}^{13+}$, $b_{232}^{12+}$) obtained from S-RBD with glycosite at Thr323.

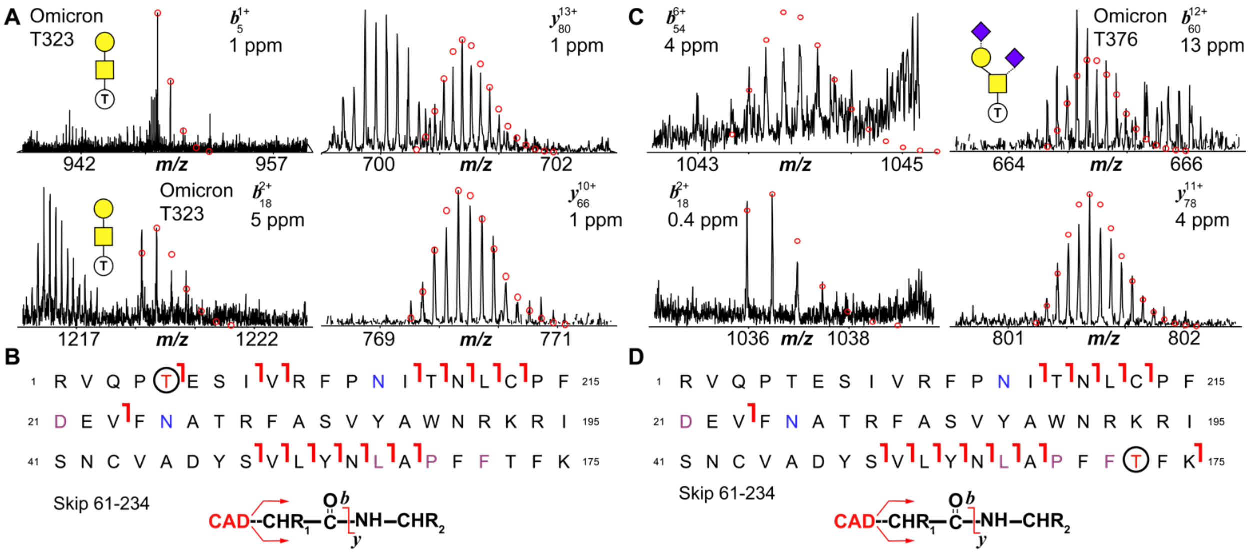


**Figure S7.** Novel T376 O-glycosylation site characterization for the Omicron S-RBD by top-down MS/MS after PNGase F treatment. (A,C) MS/MS characterization of S-RBD proteoform isolated from quadrupole window centered at 1069.4 *m/z*. Glycoform characterization reveals the specific S-RBD proteoform to have Core 1 type GalNAcGalNeuAc₂ glycan with O-glycosite (A) T323 and (C) T373. Representative CAD fragment ions ($b_5^{1+}$, $b_{18}^{11+}$, $y_{80}^{13+}$, $y_{66}^{10+}$) are shown. (B,D) Fragmentation mapping of the specific S-RBD proteoform for the (A) T323 and (C) T373 GalNAcGalNeuAc₂ identified glycan. Amino acid sequence (Arg319- Phe541) was based on the entry name P0DTC2 (SPIKE_SARS2) obtained from the UniProtKB sequence database accounting for Omicron (B.1.1.529) mutations. The blue N (Asn) denotes deamidation following PNGase F treatment. Specific Omicron residue mutations are denoted with a pink residue. Theoretical ion distributions are indicated by the red dots and mass accuracy errors are listed for each fragment ion.
